# Hsa_circ_0005397 could promote hepatocellular carcinoma progression and metastasis through EIF4A3

**DOI:** 10.1101/2023.08.30.555568

**Authors:** Liu-Xia Yuan, Mei Luo, Ruo-Yu Liu, Hui-Xuan Wang, Lin-Ling Ju, Feng Wang, Li-Ya Cao, Zhong-Cheng Wang, Lin Chen

**Affiliations:** Institute of Liver Diseases, Nantong Third People’s Hospital, Affiliated Nantong Hospital 3 of Nantong University, Nantong, Jiangsu, 226000, China; Medical School of Nantong University, Nantong Third People’s Hospital, Nantong 226000, Jiangsu, China; Medical School of Nantong University, Affiliated Hospital of Nantong University, Nantong 226000, Jiangsu, China; Preventive Health Department, Nantong Third People’s Hospital, Affiliated Nantong Hospital 3 of Nantong University, Nantong, Jiangsu, 226000, China; Hepatology Department of integrated Chinese and Western Medicine, Nantong Third People’s Hospital, Affiliated Nantong Hospital 3 of Nantong University, Nantong, Jiangsu, 226000, China

**Author notes:** Correspondence: Lin Chen, Zhong-Cheng Wang. These authors have contributed equally to this work (Liu-Xia Yuan, Mei Luo, Ruo-Yu Liu).

**Keywords:** hepatocellular carcinoma, hsa_circ_0005397, EIF4A3

## Abstract

**Purpose:** The purpose was to explore the expression and potential mechanism of hsa_circ_0005397 in hepatocellular carcinoma metabolism.

**Methods:** The quantitative real-time PCR (qPCR) was used to measure the expression of hsa_circ_0005397 and EIF4A3. The specificity of primers was confirmed by agarose gel electrophoresis. The ROC curve was draw to analysis clinical value. The actinomycin D assay and Nuclear and Cytoplasmic Extraction assay were utilized to evaluate the characteristic of hsa_circ_0005397. The CCK-8 and colony formation assays were performed to detect cell proliferation. The flow cytometry analysis was used to detect the cycle distribution. The transwell assays and Xenograft tumor model were conducted to explore cell metabolism. The RNA-binding proteins of hsa_circ_0005397 in HCC were explored in bioinformatics websites. The relationship between hsa_circ_0005397 and EIF4A3 was verified by RIP assays and rescue experiments.

**Results:** Hsa_circ_0005397 and EIF4A3 were overexpressed in HCC. Through ROC analysis, hsa_circ_0005397 shown a big role in diagnosis and prognosis. Hsa_circ_0005397 was stable and almost distributed in the cytoplasm. The upregulation of hsa_circ_0005397 generally resulted in stronger proliferative ability, clonality, metastatic potency of HCC cells, while downregulation of hsa_circ_0005397 yielded opposite results. Tumor volume and size were notably reduced while downregulation of hsa_circ_0005397, showing significant difference in tumor growth. EIF4A3 was the RNA-binding protein of hsa_circ_0005397, the expression of hsa_circ_0005397 decreased equally when depletion of EIF4A3.

Knockdown of EIF4A3 could reverse the function on HCC progression.

**Conclusions:** Hsa_circ_0005397 could promote the progression of hepatocellular carcinoma through EIF4A3. These research findings may present a novel clinical value for HCC.

## Introduction

Hepatocellular carcinoma (HCC) is the most common class of primary liver cancer all over the world (1). It is all known that viral hepatitis, alcohol abuse, and nonalcoholic hepatic steatosis are the main risk factors of HCC (2, 3). Because of the concealed incidence of HCC, most patients are already in the advanced stage at the first time of diagnosis. Despite recent achievements in HCC treatment, the diagnosis and prognosis of HCC remains challenging (4, 5). Therefore, seeking effective diagnosis and treatment is of significance currently. As reported in our previous study, plasma has_circ_0005397 represented a promising biomarker for HCC, holding promise for early HCC detection(6). However, the deeper underlying mechanisms of HCC mechanism has not been concluded. Circular RNAs (circRNAs) is kind of non-coding RNA that are formed through the backsplicing of linear RNA, resulting in a circular structure(7). Due to their stable structure, making them resistant to degradation(8). Furthermore, circRNAs widely distributed in body fluids and blood, underscoring their potential as stable biomarkers(9, 10). What’s more, circRNAs participated in cancer pathogenesis, invasion and metastasis. For example, circ_103809 could target miR-620 to suppress proliferation and invasion in HCC (11). Likewise, hsa_circ_104348 could modulate miR- 187-3p/RTKN2 axis in hepatocellular carcinoma progression (12). These numerous reports indicated that CircRNAs may be a promising biological target for HCC diagnosis and prognosis.

Recently, numerous RNA-binding proteins (RBPs) have increased substantially owing to the development of new technologies (13). RBPs could influence RNA biology through post-transcriptional, an increasing number of evidence suggests that abnormal RBP expression and function are associated with cancer metastasis(14). As reported, circRNAs could regulate the biological activity of downstream targets by binding to RBPs (15). EIF4A3 (eukaryotic translation initiation factor 4A3) is a commonly observed RBP involved in post-transcriptional regulation(16, 17). For example, EIF4A3 interacted with circ_0084615 which promoted the progression of colorectal cancer via miR-599/ONECUT2 (18). Additionally, EIF4A3 could bind with has circ_0004296, inhibiting the metastasis of prostate cancer(19). What’s more, hsa_circ_0068631 could recruit EIF4A3, which increased the expression of c-Myc in Breast cancer, showing important role in cancer metabolism(20). As reported, hsa_circ_0005397 is a kind of CircRNA originated from RHOT1, which could promote HCC by regulating miR-326/PDK2 axis(21). Despite the recent report on hsa_circ_0005397 in HCC, hsa_circ_0005397 could involve in other pathogenesis with various binding sites. In this study, we found the meaningful interaction between hsa_circ_0005397 and EIF4A3 in HCC progression and metabolism.

## Materials and methods

### Cell culture and transfection

The normal hepatocyte cell line (LO2) and HCC cell lines (SMMC-7721, SK-Hep1, HCCLM3, BEL-7404, Huh7) were acquired from the Type Culture Collection Cell Committee of the Chinese Academy of Sciences (Shanghai, China). Cells were cultured in suitable Medium containing 10% fetal bovine serum (Lonsera, USA) at 37℃ with 5% CO2.

### Construction of plasmids, siRNAs, Lentiviral Vectors

The siRNA targeting the negative control (si-NC), EIF4A3 (sh EIF4A3) and hsa_circ_0005397 (si-1, si-2) were obtained from Genepharma Co., Ltd. The virus vectors were purchased from Gechem (Shanghai, China). We use Lipofectamine3000 and P3000 (Invitrogen, USA) for transfection under the guidance of manufacturer’s instruction.

### RNA preparation and qPCR

Total RNA was isolated by Trizol reagent (Invitrogen, USA), after reversing RNA into cDNA (Thermo, USA). SYBR GreenⅠ Master (Roche, German) was used for qPCR. All data were normalized by 18S and the relative expression were calculated by 2–ΔΔCT method. The hsa_circ_0005397 primers were purchased from Geneseed (Guangzhou, China), the other primers were purchased from Sangon Biotech (shanghai, China). Primers used in this study are as follows: hsa_circ_0005397 (Forward: 5′- GACAAAGACAGCA GGTTCCT-3′; Reverse: 5′-CTCTGTTCTGCTTCT GAGTA-3′); EIF4A3 (Forward: 5’-CAGGGCGTGTTTTTGATATGAT-3’; Reverse: 5’-ATCAGCTTCATCCAAAACCAAC-3’; RHOT1(Forward: 5’- CTGATTTCTGCAGGAAACACAA-3’; Reverse: 5’- GCAAAAACAGTAGCACCAAAAC-3’).18S rRNA (Forward: 5’- GTAACCCGTT GAACCCCATT-3’; Reverse: 5’- CCATCCAATCGGTAGTAGCG-3). After qPCR, the PCR products were added to 2% agarose gel to electrophoresis.

### RNase R resistance assay

RNase R (Baiye Biotechnology Center, China) treatment was utilized to degrade linear RNA. Briefly, total RNA was incubated with RNase R or without RNase R for 0.5 h at 37℃. After that, the treated RNA was used for qRT-PCR to examine the levels of hsa_circ_0005397 and RHOT1.

### Nuclear and Cytoplasmic Extraction

Total RNA was separated by nuclear and cytoplasmic protein extraction kits under the construction (Beyotime, China).

### Actinomycin D assay

2.5mg/mL Actinomycin D (Merck, German) was added into cell cultured until 24 hours. Trizol Reagent (Invitrogen, USA) was added at 0, 2, 4, 6, 8 hours to collect RNA.

### Cell Counting kit-8 (CCK-8) assay

After 48 hours, the transfected cells were collected and 3,000 cells/well were seeded in 96-well plate. While incubated with CCK-8 solution (MCE, USA) for 2 hours, the OD value was measured at 450 nm (Thermo,USA).

### Colony formation assay

3,000 cells were added into a six-well plate until two weeks, fixed with 4% formaldehyde (Beyotime, China), stained with 0.1% crystal violet (Sigma, USA) and photographed finally.

### Transwell assay

The 8μm pore size chamber (Costar, USA) and Matrigel (BD, USA) were used for cell migration and invasion assays. After transfected 48 hours, 5x10^4cells were suspended in serum-free media into the upper chambers, while the lower chambers were added with media containing 20% FBS in the migration assay. Similarly, part of transfected cells was added in upper chambers which precoated with 100ul Matrigel for invasion assay. Cells were fixed with 4% paraformaldehyde, stained with 0.1% crystal violet (Sigma, USA) and photographed under a microscope (Olympus, Japan).

### Website Prediction

We use Circ Interactome (https://circinteractome.nia.nih.gov/) and CircFunBase (https://bis.zju.edu.cn/CircFunBase/index.php) to search the RNA-binding Proteins matched to has_circ_0005397, Platform could contribute to CircRNA-Protein studies(22, 23).

### Flow cytometry analysis

The transfected cells were collected and fixed in ice-cold ethanol overnight. Under the constructions, we added 500μL propidium iodide (PI) staining solution (Solarbio, Beijing) into each sample, incubating 30 min at 37℃, then cooling at 4℃, the cell cycle difference was estimated by flow cytometer (BIORAD, USA).

### RNA immunoprecipitation (RIP) assay

The experiment was conducted with RIP Kit (Geneseed, Guangzhou). The cells were suspended in Buffer A contained with inhibitors to collect the whole lysate. The lysate was incubated by RIP beads against human anti- Argonaute2 (Anti-Ago2) or anti-immunoglobulin G (Anti-IgG) antibody at 4℃, the purified RNA was extracted and detected by qPCR.

### Xenograft tumor assay

HCCLM3 cells transfected with pcDNA5.1-circ_0005397 or pcDNA5.1-NC were injected into the subcutaneous of nude mice (n = 3). The volume and size were checked every day until all mice sacrificed, the tumors were collected to estimate the efficiency of has_circ_0005397. All animal experiments were complied with the rules of the Ethics Committee of Affiliated Nantong Hospital 3 of Nantong University.

### Statistical analysis

Through at least three independent experiments, the data were presented as the mean ± standard deviation (SD) and analyzed by GraphPad Prism 7.0 (GraphPad Software, USA). The t-test and one-way analysis was employed for analyzing differences between two groups or more than two groups were compared. The correlation was calculated by Pearson’s correlation analyses. P value <0.05 were considered significant.

## Results

### The characteristics of hsa_circ_0005397 in Hepatocellular Carcinoma

Firstly, hsa_circ_0005397 was upregulated in HCC tissues (Fig 1A) andhigher hsa_circ_0005397 was associated with TNM stage and tumor size (**Table1)**. According to the ROC analysis, the area under the ROC curve was 0.8763 (95% CI 0.81-0.93), with a sensitivity of 84.21% (95% CI 0.72-0.91) and a specificity of 82.46% (95% CI 0.71-0.90) (Fig 1B). At the same time, HCC cell lines presented higher expression compared with LO2 cell (Fig 1C), and the primers and product were specificity (Fig 1D). We confirmed that hsa_circ_0005397 could resist the digestion of RNase R, suggesting the cyclic structure of hsa_circ_0005397(Fig 1E). Furthermore, we utilized the nuclear and cytoplasmic extraction kits to explored the distribution of hsa_circ_0005397, indicated that hsa_circ_0005397 located in the cytoplasm mostly (Fig 1F). Finally, the actinomycin D experiments proved the longer half-time of hsa_circ_0005397, showing a better stability (Fig 1G).

**Table 1.**
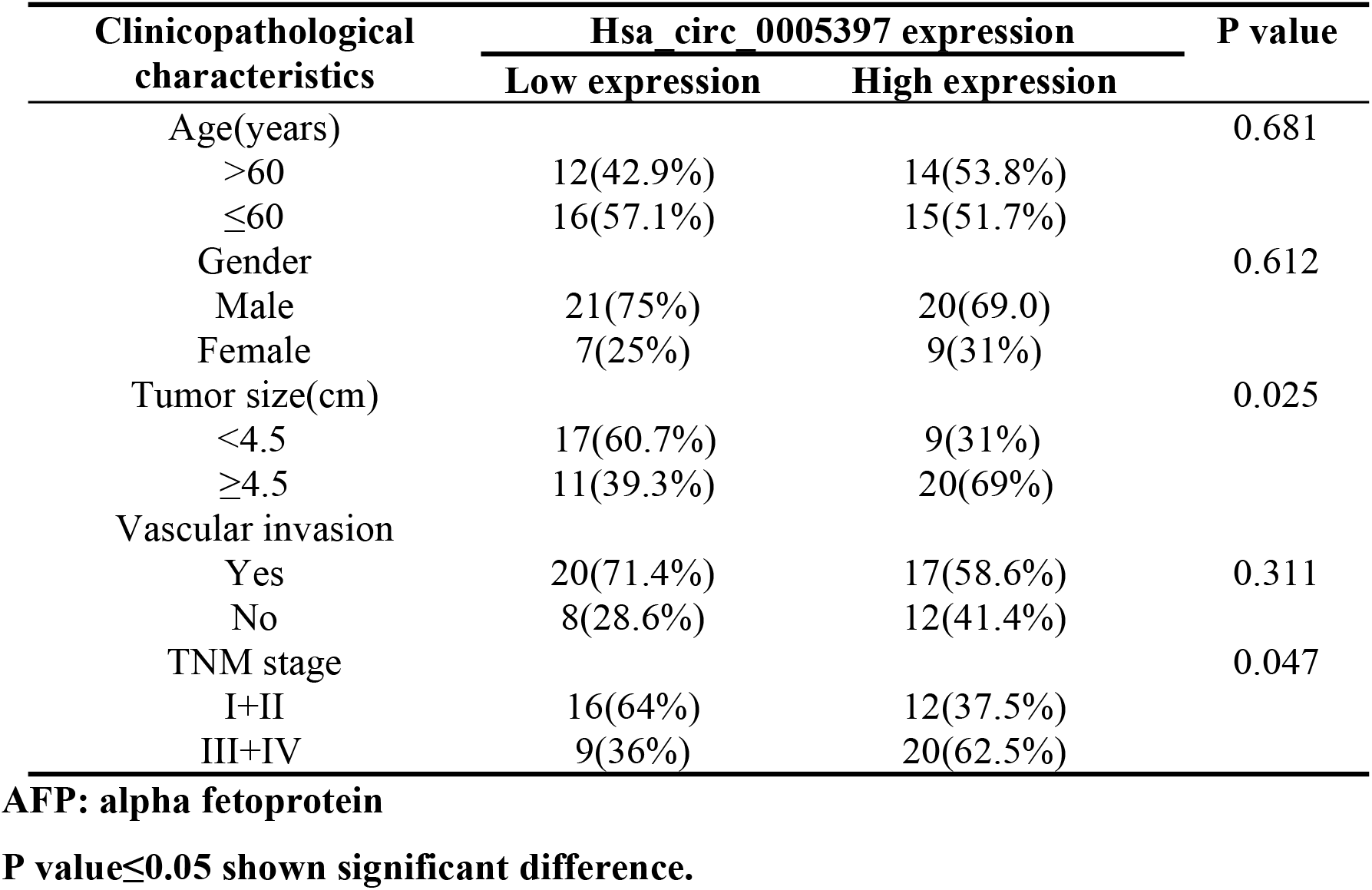
Relationship between Clinicopathological Characteristics and hsa_0005397 expression in patients.

**Fig 1.**
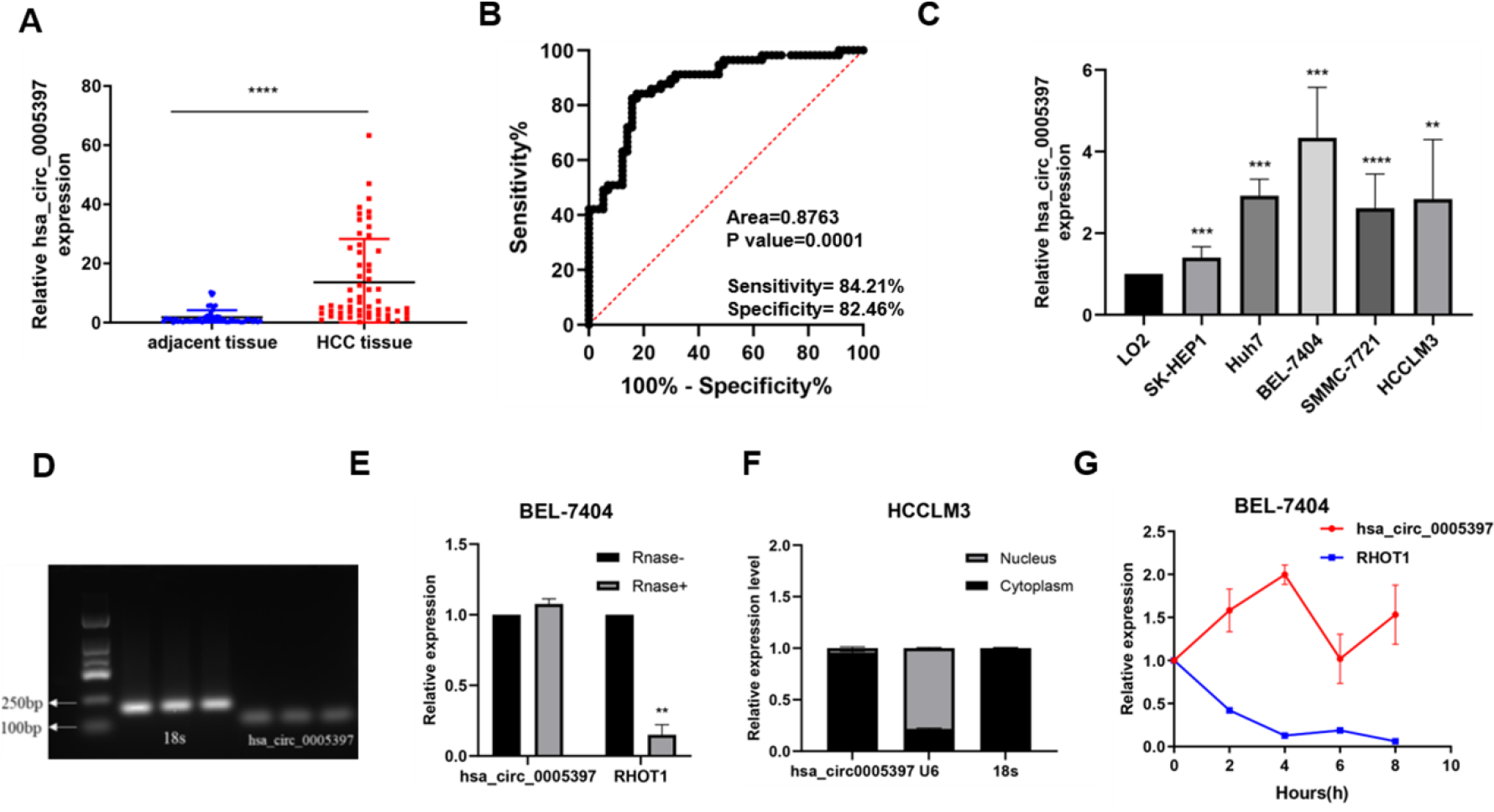
The characteristics of hsa_circ_0005397 in Hepatocellular Carcinoma. (A)The expression of hsa_circ_0005397 in HCC tissues; (B) The ROC curve of hsa_circ_0005397 (C) The expression of hsa_circ_0005397 in cell lines; (D) The PCR products were performed by gel electrophoresis; (E) The relative abundance after treated with or without RNase R in BEL-7404; (F)The distribution of hsa_circ_0005397; (G) qPCR for expression of hsa_circ_0005397 and RHOT1 mRNA in BEL-7404 cells treated with Actinomycin D at different point. *P<0.05, **p<0.01, ***p<0.001.

### Hsa_circ_0005397 could affect proliferation in HCC cell lines

QPCR analysis revealed that si-hsa_circ_0005397 (si1, si2) significantly silenced the expression of hsa_circ_0005397 In HCCLM3 and BEL-7404 cells, while the high efficiency of overexpression in SK-Hep1 (Fig 2A). The CCK-8 assays showed that overexpression of hsa_circ_0005397 could promote cell proliferation, consistently silenced of hsa_circ_0005397 inhibited the cell proliferation (Fig 2B). Meanwhile, compared with the negative control, the hsa_circ_0005397-overexpression group raised the number of clones in SK-Hep1 cells, in contrast to the hsa_circ_0005397- depletion group (Fig 2C-D).

**Fig 2.**
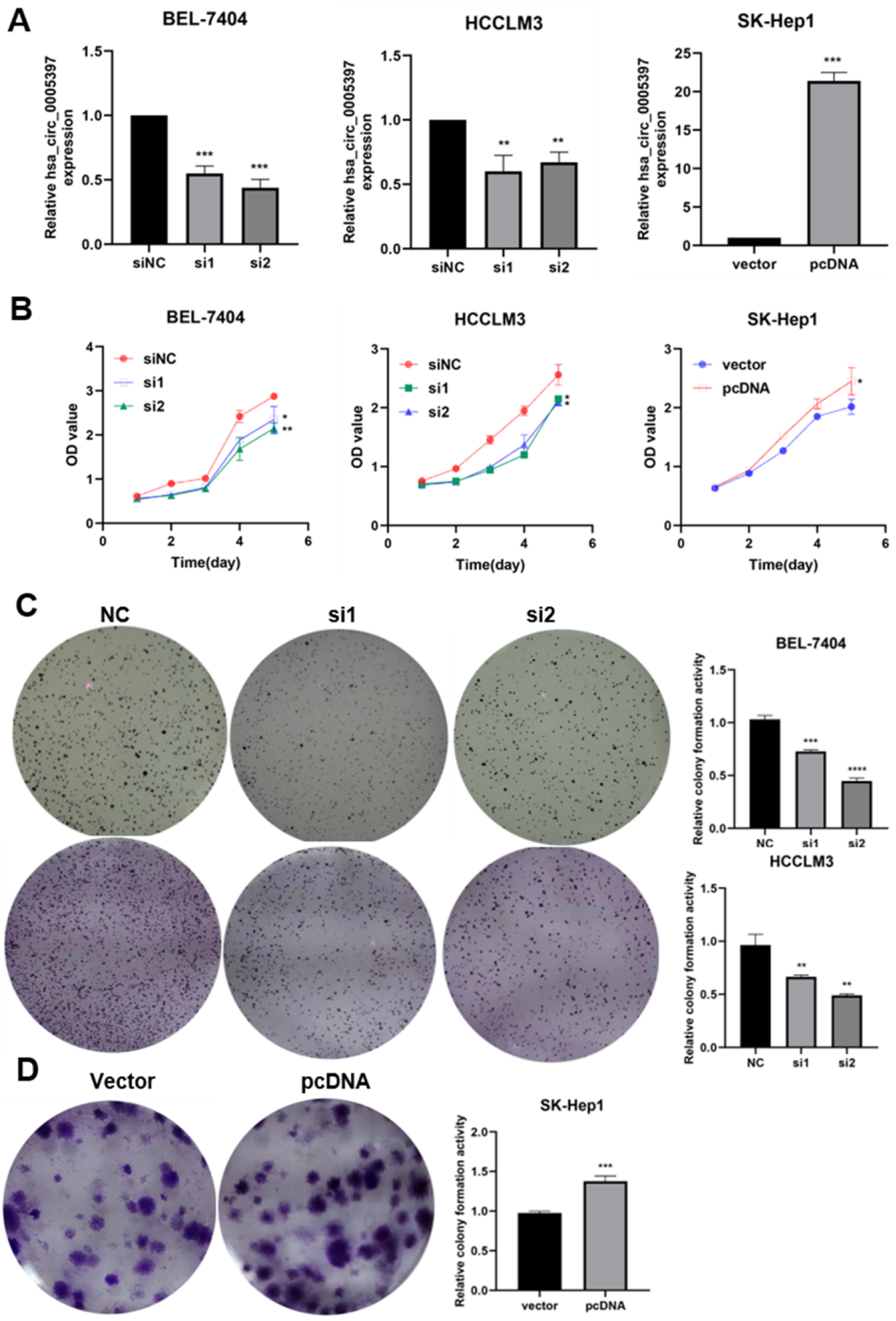
Hsa_circ_0005397 could affect proliferation in HCC cell lines. (A)The expression of hsa_circ_0005397 in BEL-7404, HCCLM3 and SK- Hep1; (B)- Cell proliferation was determined by CCK-8 assay (D) colony formation was photographed and analyzed by Image J. *P<0.05, **p<0.01, ***p<0.001.

### Hsa_circ_0005397 could affect migration and invasion capacity in HCC cell lines

Furthermore, we explored the effects of hsa_circ_0005397 on migration and invasion in HCC cell lines by transwell assays, the relative migration and invasion (Fig 3A-C) were increased in the hsa_circ_0005397-overexpression group, while decreased in the hsa_circ_0005397-depletion group, showing a significant difference in migration and invasion.

**Fig 3.**
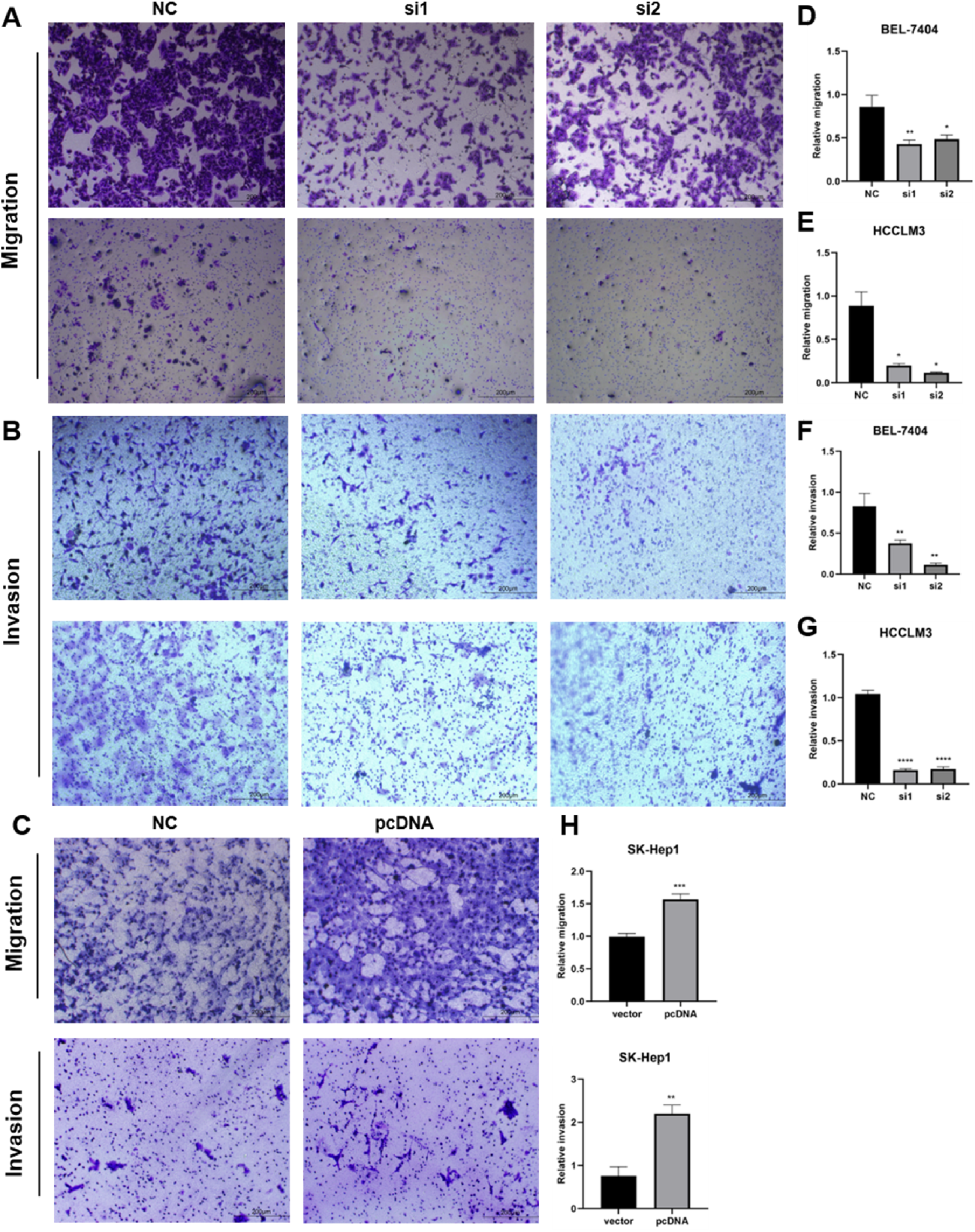
Hsa_circ_0005397 could affect migration and invasion in HCC cells. (A-C) Transwell assays were used to evaluate cell migration and invasion ability in BEL-7404 and HCCLM3 transfected with NC, si1, si2, while overexpression of hsa_circ_0005397 in SK-Hep1 with vector and pcDNA ((Magnification,20X; Scale,200μm); (D-H) The relative migration and invasion percentage were evaluated by ImageJ. *P<0.05, **p<0.01, ***p<0.001.

### Hsa_circ_0005397 could affect cell cycle in HCC cell lines

Furthermore, cell cycle was tested, indicating that the percentage of the G0/G1phase increased in hsa_circ_0005397 depletion group (Fig 4A-B), while decreased in hsa_circ_0005397-overexpression group (Fig 4C). The percentage of cell phase was shown in Fig D-F. All results above proved that hsa_circ_0005397 promotes the growth in HCC cell lines.

**Fig 4.**
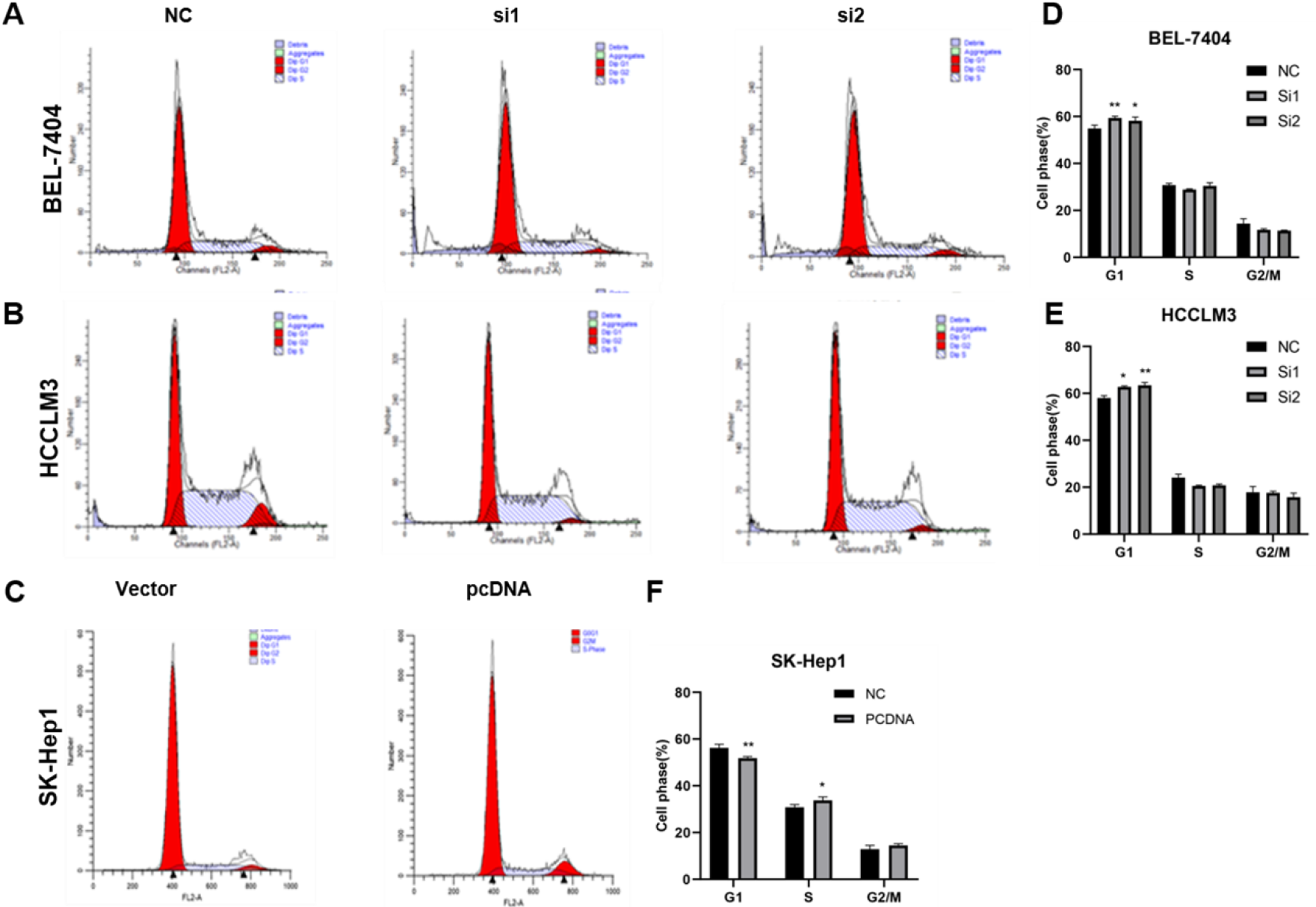
Hsa_circ_0005397 effect cell cycle in HCC cells. (A-C) The represented pictures in cell cycle; (D-E) The cell phase (%) were analyzed by Graphpad Prism. *P<0.05, **p<0.01, ***p<0.001.

### Hsa_circ_0005397 could promote tumor growth in vivo

In order to explore the role of hsa_circ_0005397 in HCC metabolism, we prepared the subcutaneous tumor formation experiments. qPCR confirmed the efficiency of virus infection in HCCLM3 cells (Fig5A). Compared with LV-shNC group, the size and weight of tumor was notably reduced while knockdown of the hsa_circ_0005397, showing the importance of hsa_circ_0005397 in tumor growth (Fig 5B-C).

**Fig 5.**
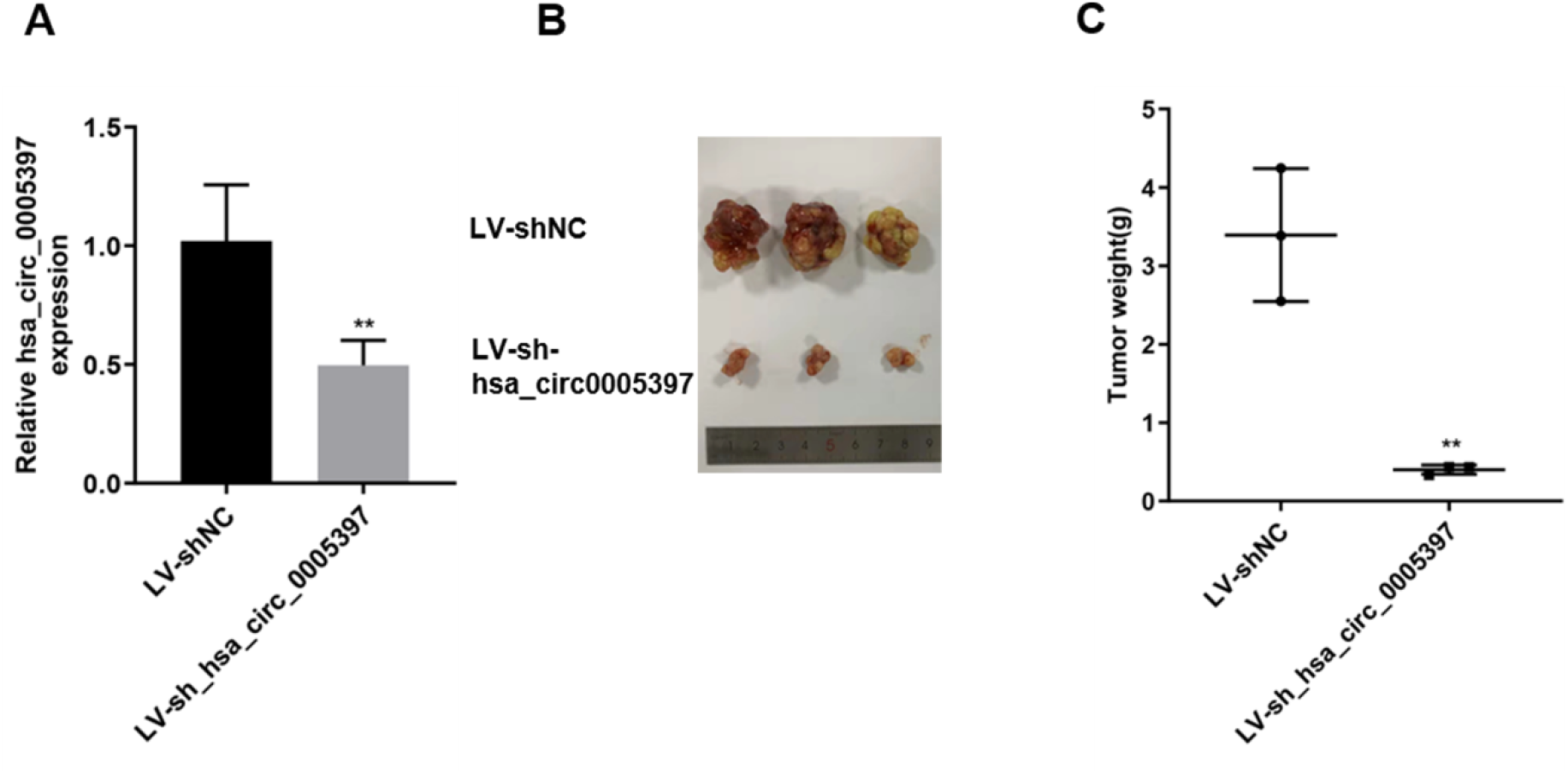
Hsa_circ_0005397 effect tumor growth in vivo. (A) The efficiency of virus infection is determined by qPCR; (B-C) The tumor size and weight were determined and analyzed by Graphpad Prism. *P<0.05, **p<0.01, ***p<0.001.

### Hsa_circ_0005397 could promote the metabolism on HCC cell lines through EIF4A3

According to the prediction of Circ Interactome and CircFunBase, EIF4A3 matched to hsa_circ_0005397. Next, we also found that EIF4A3 was overexpressed in HCC cell lines and tissues (Fig 6B-C). Moreover, the expression of EIF4A3 and hsa_circ_0005397 has a significant positive correlation (Fig 6D). The efficiency of EIF4A3 depletion is verified by qPCR (Fig 6E). After depletion of EIF4A3, the expression of hsa_circ_0005397 decreased significantly in HCCLM3 (Fig 6F). We used RIP assay to verify the relationship between hsa_circ_0005397 and EIF4A3. Compared to the negative control IgG group, hsa_circ_0005397 was enriched in the EIF4A3 group (Fig 6G). Next, we designed rescue experiments, HCCLM3 cells were co-transfected with pcDNA-circ_05397 and shEIF4A3, qPCR was used to verify the expression of circ_0005397 (Fig 6H). Meanwhile, the functional experiments showed that overexpression of hsa_circ_0005397 could promote the proliferation, migration and invasion ability of HCC cells, while knockdown of EIF4A3 could lead to the opposite results (Fig 6I-J). Consistently with these findings, EIF4A3 remarkably regulated HCC cell migration and proliferation in HCC cells. suggesting that a unique relationship between EIF4A3 and hsa_circ_0005397. In conclusion, these results demonstrated that hsa_circ_0005397 could promote the proliferation and metabolism of HCC through EIF4A3.

**Fig 6.**
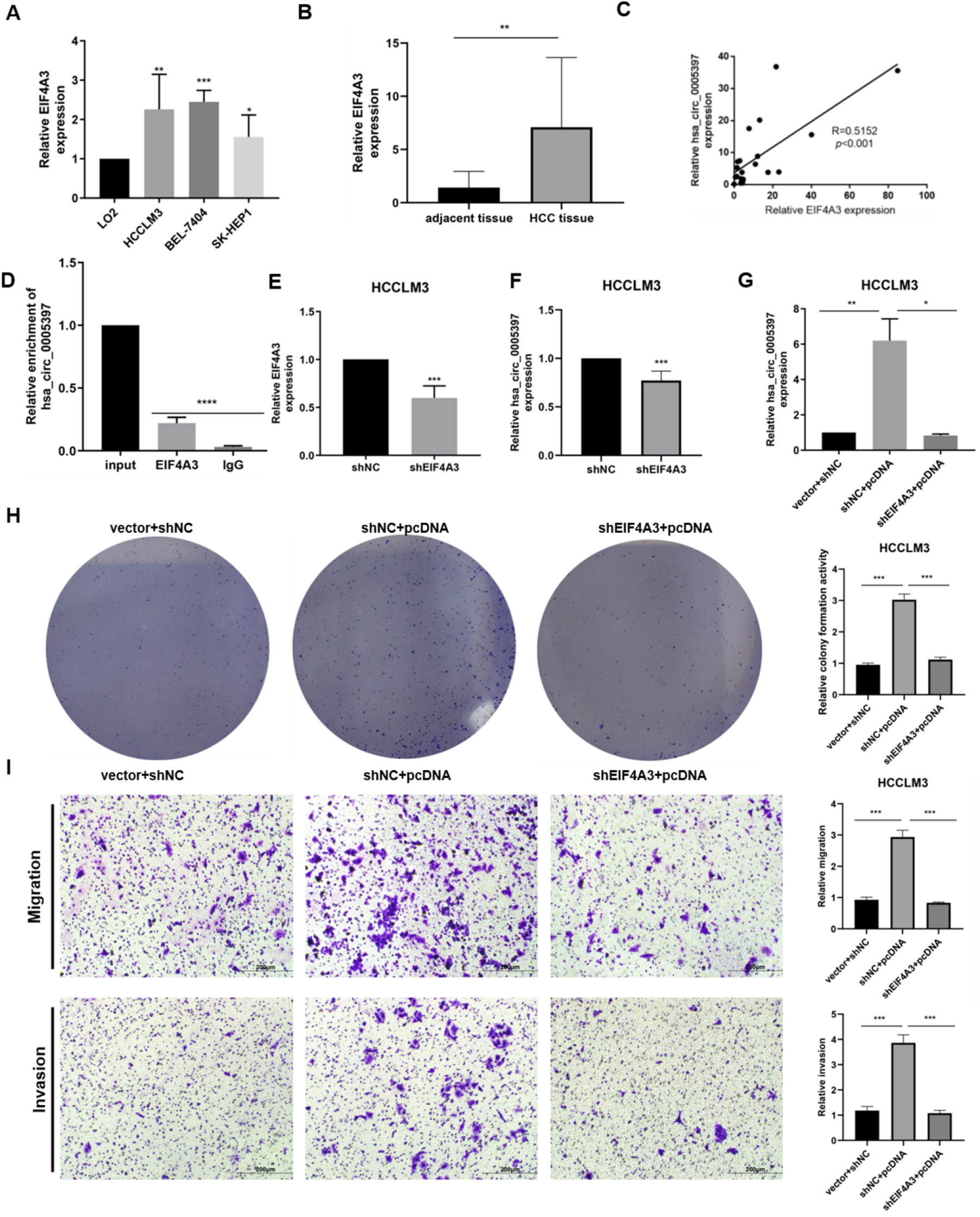
Hsa_circ_0005397 could promote the metabolism on HCC cell lines through EIF4A3. (A) The EIF4A3 expression in HCC cell lines; (B) The EIF4A3 expression in HCC tissues; (C) The relationship between EIF4A3 and hsa_circ_0005397; (D) RIP assay to verify the relationship between hsa_circ_0005397 and EIF4A3; (E) The expression in HCCLM3 transfected with shNC or shEIF4A3; (F) The expression of hsa_circ_0005397 in HCCLM3 while transfected with shNC or shEIF4A3; (G) The expression of hsa_circ_0005397 in HCCLM3 transfected with shNC+vector, shEIF4A3+pcDNA or shEIF4A3+pcDNA; (H) Representative images of clone formation analysis and the relative colony ability was evaluated by Image J; (I) Cell migration and invasion observed in transwell assay (magnification,20x; Scale bar,200μm) and the relative migration and invasion is counted by Image J and analyzed by Graphpad Prism. *P<0.05, **p<0.01, ***p<0.001.

## Discussion

Although advancements have been made in this field (24, 25), due to challenges in early tumor diagnose, as well as tumor recurrence and metastasis the recent survival rate of patients with HCC remains unsatisfactory (26, 27). Although serum AFP has been widely utilized as a tumor marker for HCC diagnosis and screening, its sensitivity and specificity are not sufficiently high, necessitating further improvements (28, 29). In recent years, research on Circular RNAs (circRNAs) has provided a novel avenue that is expected to contribute to the clinical diagnosis and prognosis of HCC(30-32).

With the development of sequencing and other technologies, the diversity and biological functions of circRNAs are being extensively explored. As reported, circRNAs could through sponging miRNAs, binding to RBPs, and regulating gene transcription and translation to regulate tumorigenesis (20, 33, 34). EIF4A3 plays a crucial role in post-transcriptional regulation, which has been reported to interact with RNAs and act as a diagnostic marker in many cancers (35-37). Based on previous reports, studies have demonstrated that circ_0004296 can sequester EIF4A3, leading to the inhibition of ETS1, thus impeding prostate cancer metastasis(19). What’s more, EIF4A3 could bind to upstream region to promote circMMP9 expression, showing the important role in mRNA splicing(38).

Recently, we have confirmed the upregulation of hsa_circ_0005397 in HCC serum, which could act as novel biomarker(6). However, the mechanism of abnormal expression in HCC tissues and cell lines has not been defined. In this study, we verified that hsa_circ_0005397 and EIF4A3 was remarkably overexpression in HCC. Functionally, hsa_circ_0005397 knockdown could inhibit HCC cell proliferation and metastasis. Also, hsa_circ_0005397 and EIF4A3 has positive correlation. Consistently, the RIP assay was performed to confirmed that EIF4A3 could interact with hsa_circ_0005397. What’s more, EIF4A3 inhibition reversed the repressive effects of si- hsa_circ_0005397 on HCC cell proliferation and invasion. All in all, these findings concluded that hsa_circ_0005397 could promote HCC progression and metastasis through EIF4A3.

## Conclusion

Our study concluded that overexpression of hsa_circ_0005397 could promote the progression and metabolism of HCC through EIF4A3.

Regulating the hsa_circ_0005397/EIF4A3 axis might be a potential strategy for HCC.

## Acknowledgments

This work was supported by Nantong Science and Technology Bureau (JCZ2022036, JCZ2022080), Jiangsu Health and Health Committee (Z2020011), Nantong Municipal Health Commission (QN2022041, QA20210039).

## Ethics approval and consent to participate

This study was approved by the Ethics Committee of Nantong Third Affiliated Hospital of Nantong University. All participants signed informed consent forms in this study.

### Conflicts of Interest Statement

No potential conflicts of interest were disclosed.

### Authors’ Contributions

**Conception and design**: Lin Chen and Zhong-Cheng Wang;

### Experiment and data acquisition

Liu-Xia Yuan, Mei Luo, Ruo-Yu Liu;

### Technical, or material support

Hui-Xuan Wang, Lin-Ling Ju, Feng Wang;

### Clinical information and patients

Mei Luo, Ruo-Yu Liu, Li-Ya Cao;

### Statistical analysis

Liu-Xia Yuan, Mei Luo, Ruo-Yu Liu;

### Writing, review, and/or revision of the manuscript

Liu-Xia Yuan, Mei Luo, Ruo-Yu Liu, Hui-Xuan Wang, Lin-Ling Ju, Feng Wang, Li-Ya Cao, Zhong-Cheng Wang and Lin Chen.

